# The Microbiomes of Pancreatic Tissue in Pancreatic Cancer and Non-Cancer Subjects

**DOI:** 10.1101/189043

**Authors:** Erika del Castillo, Richard Meier, Devin C. Koestler, Tsute Chen, Bruce J. Paster, Kevin P. Charpentier, Karl T. Kelsey, Jacques Izard, Dominique S. Michaud

## Abstract

**Objective:** To determine whether bacteria are present in the pancreas of pancreatic cancer and non-cancer subjects and examine whether bacterial profiles vary by site and disease phenotype.

**Design:** 77 patients requiring surgery for pancreatic diseases, or diseases of the foregut, at the Rhode Island Hospital (RIH) were recruited into this study between 2014 and 2016. In addition, 36 whole pancreas were obtained from the National Disease Research Interchange (NDRI) from subjects who were of similar age as the RIH patients and had not died of cancer. The primary exposure of interest was the measurement of the relative abundance of bacterial taxa in all tissue specimens using 16S rRNA gene sequencing.

**Results:** Number of bacterial reads per sample varied substantially across sample type and patients, but all demonstrated the presence of diverse gastrointestinal bacteria, including bacterial taxa typically identified in the oral cavity. Bacterial profiles were noted to be more similar within individuals across sites in the pancreas, than between individuals by site, suggesting that the pancreas as a whole has its own microbiome. Comparing the mean relative abundance of bacterial taxa in pancreatic cancer patients to those without cancer revealed differences in bacterial taxa previously linked to periodontal disease, including *Porphyromonas*.

**Conclusions:** Bacterial taxa known to inhabit the oral cavity, as well as the intestine, were identified in pancreatic tissue of cancer and non-cancer subjects. Whether any of these bacteria play a causal role in pancreatic carcinogenesis, or are simply opportunistic in nature, needs to be further examined.

## Introduction

In 2017, an estimated 53,670 individuals will be diagnosed with pancreatic cancer in the US, and only 9% of these individuals are expected to survive the next five years ^1^. Given this high fatality rate, and the silent progression of early disease, identifying risk factors for the prevention and early detection of pancreatic cancer is critical to reducing its mortality. To date, known risk factors for pancreatic cancer, including smoking, obesity, diabetes, heavy alcohol consumption, family history and markers of genetic susceptibility, cannot, even collectively, be used for early detection and risk stratification of pancreatic cancer in the general population ^2^.

Studies have suggested a link between bacteria and pancreatic cancer risk ^3^, highlighting the need to explore the currently unknown role of bacteria in the diseased pancreas. The current research on oral bacteria and pancreatic cancer risk stems from a number of observational studies that reported a higher risk of pancreatic cancer among individuals with periodontitis, when compared to those without periodontitis ^3^ ^4^. Periodontitis, an inflammatory disease of the gums, is largely driven by dysbiosis promoting pathogenic oral bacteria ^5^. Two large prospective cohort studies have reported positive associations between periodontal disease pathogens and subsequent pancreatic cancer risk ^6^ ^7^; in these two studies, detection of elevated antibodies to *Porphyromonas gingivalis*, measured in blood collected prior to cancer diagnosis, was associated with a two-fold higher risk of pancreatic cancer ^6^, and presence (vs absence) of *Porphyromonas gingivalis* in saliva collected prior to cancer diagnosis was associated with a 60% increase in risk of pancreatic cancer ^7^. *Aggregatibacter actinomycetemcomitans*, another periodontal pathogen, was also associated with pancreatic cancer risk in the prospective study using saliva ^7^.

Few investigations to date have attempted to detect bacteria in, or around, the pancreas. Earlier studies reported the presence of bacteria in pancreatic ducts of patients with chronic pancreatitis or bile duct obstruction ^8-10^. Other studies have investigated the presence of specific bacterial DNA in the pancreatic tissue of pancreatic cancer patients, namely species of *Helicobacter* ^11^ and *Fusobacterium* ^12^. The most comprehensive microbiome study to date reported the presence of a diverse bacterial population in fluid collected from the bile duct, pancreas and jejunum of patients undergoing pancreaticoduodenectomy ^13^.

Metagenomics studies on DNA isolated from tissue samples from cancer patients have been conducted for lung ^14^, esophageal ^15^, stomach ^16^, and breast cancer ^17^. These studies demonstrate that 16S rRNA gene sequencing can be effectively conducted on fresh tissue samples, where the ratio of bacterial to human DNA is much lower than at other human sites (e.g., stool or oral cavity)^18^. Moreover, these studies have shown that bacterial profiles at different organ sites are often unique ^14 17^ and that changes may be associated with cancer ^15 16^.

To date, no study has identified the overall microbiome in pancreatic and surrounding tissues samples in normal and diseased individuals, a critical step to understand whether and how bacteria may play a role in carcinogenesis. In an effort to address the specific question of whether the pancreas has its own microbiome in cancer patients, we recruited patients from the Rhode Island Hospital (Providence, RI) with planned foregut surgery to obtain tissue samples for 16S rRNA gene microbiome analysis. In addition, we obtained non-diseased pancreatic tissue from National Disease Research Interchange (NDRI) for comparison.

## Materials and Methods

### Study population and sample collection

A total of 77 subjects, enrolled between January 2014 and March 2016, were included in this study. Subjects were eligible if identified as candidates for surgery of the foregut by Dr. Charpentier, the lead surgeon at the RIH; patients included those with pancreatic cancer, pancreatic cystic neoplasms, pancreatitis, bile duct or small bowel diseases. All the recruited patients were between 31-86 years old (Table 1). Participants were asked to complete a self-administered questionnaire to provide data on demographic and behavioral factors. Questions on family history of cancer, use of antibiotics and other over the counter medications were also included on the questionnaire. Stool collection kits with ethanol as a fixative were provided prior to surgery ^19^; participants were asked to return the samples using a pre-paid box.

**Table 1.** Distribution of demographic, lifestyle and health conditions variables among patients with diseases of the foregut, primarily pancreatic diseases, and deceased controls.

| RIH Subjects (n=77) |  | NDRI Subjects (n=36) |  |
| --- | --- | --- | --- |
| Characteristic | Mean (SD) | Characteristic | Mean (SD) |
| Age | 63 ± 13 | Age | 69 ± 15 |
| BMI | 27 ± 6 | BMI | 29 ± 6 |
| N (%) |  | N (%) |  |
| <u>Sex</u> |  | <u>Sex</u> |  |
| Male | 38 (49) | Male | 22 (61) |
| Female | 39 (51) | Female | 14 (39) |
| <u>Race</u> |  | <u>Race</u> |  |
| Caucasian | 72 (93.5) | Caucasian | 32 (88.9) |
| Black | 2 (2.6) | Black | 2 (5.6) |
| Other | 2 (2.6) | Other | 2 (5.6) |
| <u>Smoking status</u> |  | <u>Smoking status</u> |  |
| Ever smoker | 44 (58) | Ever smoker | 24 (67) |
| <u>Chemotherapy</u> |  | <u>Cause of Death</u> |  |
| Never | 52 (76.5) | Heart failure | 17 (47.2) |
| Prior to past 6 months | 7 (10.3) | Cardiopulmonary arrest | 6 (16.7) |
| In past 6 months | 9 (13.2) | Cerebrovascular accident | 2 (5.6) |
| <u>Antibiotic use</u> |  | Respiratory arrest | 2 (5.6) |
| Never | 13 (18.1) | Abdominal aortic aneurysm | 1 (2.8) |
| Prior to past 6 months | 32 (44.2) | Intracerebral hemorrhage | 1 (2.8) |
| In past 6 months | 21 (29.2) | Liver cirrhosis | 1 (2.8) |
| Missing | 6 (8.3) | Overdose | 1 (2.8) |
| Stent prior to surgery (yes) | 19 | Parkinson's disease | 1 (2.8) |
| Preop EUS | 20 | Pneumonia | 1 (2.8) |
| <u>Surgery for:</u> |  | Pulmonary embolism | 1 (2.8) |
| Pancreatic cancer | 51 (66.2) | Pulmonary fibrosis | 1 (2.8) |
| Chronic pancreatitis | 16 (20.8) |  |  |
| Other | 10 (13.0) |  |  |

A protocol was established for processing tissue samples collected during surgery to reduce contamination. A technician from the Pathology Department was informed in advance of the surgery date and time, and was paged once the specimens were obtained. Surgical tissue samples were frozen within one hour of the surgery time, as well as tissues swab samples from the stomach, jejunum, and bile duct that were collected using DNA-free forensic sterile swabs whenever possible. During surgery, the surgeon also recorded (on a surgery form for the study) if the patient had received prior pre-OP EUS, had previously had their gallbladder removed, and had received prior placement of a stent (for treatment of symptoms); all patients received one dose of perioperative antibiotics prior to surgery. Tissue samples (pancreatic tumors, pancreatic cysts, normal pancreas, pancreatic ducts and duodenums) were prepared by a Rhode Island Hospital pathologist; cancerous and non-cancerous tissues were identified, separated and labeled. All samples were de-identified and stored at -80°C until processing.

Upon review of pathology records, ICD10 codes were assigned to each subject; 39 subjects had pancreatic cancer (ICD10 codes C25.0-C25.9), 12 subjects had periampullary cancer (ICD10 codes C24.0-C24.1), 18 subjects had other pancreatic conditions (ICD10 codes K86.0-K86.3), and the remaining 8 had other gastrointestinal conditions. The study was approved by Lifespan’s Research Protection Office for recruitment at RIH, as well as the Institutional Review Boards for human subjects research at Brown University, Tufts University and Forsyth Institute.

In addition, we obtained pancreatic specimens without known conditions of pancreatic diseases from the National Disease Research Interchange (NDRI) to serve as control samples. Snap-frozen ‘control’ whole-pancreas and duodenum (∼ 5cm) human specimens from 36 deceased donors were obtained from NDRI with an average post-mortem recovery time of 13.6 hours. Control pancreas (head and tail), pancreatic ducts and duodenums were dissected under sterile conditions, and stored at -80°C until processing. To remove additional contamination, we removed a thin tissue layer around each sample prior to extracting DNA.

Details for DNA extraction and sequencing procedures are provided in the Supplementary Methods.

### 16S rRNA amplicon Illumina sequencing

The 16S rRNA gene dataset consists of Illumina MiSeq sequences targeting the V3-V4 hypervariable regions. The DNA target sequencing was performed by the Forsyth Institute Sequencing Core. To validate batch sequencing, bacterial mock community samples were used; their relative abundance were consistent across the MiSeq runs.

The MiSeq reporter analysis was used to discard low quality sequences and to generate FASTQ files containing only filtered quality sequences, subsequently the overlapping paired-end reads were stitched together and further processed using used a multi-stage BLASTN-base search taxonomy read assignment pipeline that maximizes species level classification ^20^.

### Taxonomic assignment pipeline of 16S rRNA amplicon sequencing data

Sequences were BLASTN-searched against a combined set of 16S rRNA reference sequences that consist of the HOMD (version 14.5), Greengenes Gold ^21^, and the NCBI 16S rRNA reference sequence set. Representative reads from each of the Operational Taxonomic Units (OTUs)/species were BLASTN-searched against the same reference sequence set again to determine the closest species for these potential novel species. All assigned reads were subject to several down-stream bioinformatics analyses, including alpha and beta diversity assessments, provided in the QIIME (Quantitative Insights Into Microbial Ecology ^22^) software package version 1.9.1.

### Statistical analysis

Samples with < 500 read counts were excluded from all analysis. In addition, only OTUs with a minimal read count of 100 sequences (across all samples) were included in the analyses. For QIIME analyses, we normalized the number of sequences in the different MiSeq runs by rarefying each library to 500 reads to account for differences in sequencing depth across runs. Across samples, OTU relative abundance was computed as the ratio of an OTU’s absolute abundance to the total number of reads for that sample.

To examine the variation in the microbial profile across the different habitats/sites (Supplemental Table 1) among the NDRI and RIH subjects, we calculated the distance/dissimilarity between samples using the Bray-Curtis and Sorenson indices ^23^. Computed distances were subsequently used to generate principal coordinate analysis (PCoA) plots to visualize the arrangement of the samples in the ordination space.

To identify demographic and clinical correlates of pancreatic microbial composition, we fit a series of zero-inflated beta regression models to examine associations between genus-level relative abundances and demographic (i.e., age, gender, race, BMI) and clinical (i.e., health status, chemotherapy, antibiotics use prior to surgery, anxiety medications, presence of stent prior to surgery, preoperative endoscopic ultrasound [Preop EUS], tumor surgery classification by International Code of Disease [ICD code]). More details are provided in the Supplemental Methods.

## Results

The present analysis included a total of 263 pancreatic tissue and swab samples collected from 113 subjects (77 patients from RIH providing 150 samples and 36 subjects from NDRI providing 113 samples; Supplemental Table 1). There were no significant differences in the distribution of age, gender, BMI, race, and smoking status between RIH and NDRI patients (Table 1). The Illumina-based sequencing of V3-V4 hypervariable regions of the bacterial 16S rRNA gene resulted in a total of 19,498,743 high quality sequences (with a median sequence length of 427 nucleotides).

### Taxonomy

Over 99% of the reads from RIH pancreatic samples were attributed to 5 bacterial phyla (45.9% Proteobacteria, 35.6% Firmicutes, 9.5 % Bacteroidetes, 4.3% Fusobacteria, and 3.9% Actinobacteria; Figure 1A). The remaining low abundance phylotypes (0.6% of the total) belonged to six bacterial phyla (Synergistetes, TM7, Deinococcus-Thermus, Verrucomicrobia, Spirochaetes, and Tenericutes). Although 99.6% of the reads observed among the NDRI pancreatic samples belonged to the same five bacterial phyla as observed in RIH patients, there were notable differences in proportions of these five phyla between the RIH and NDRI samples (Figure 1A). The phylum Tenericutes (Bacteria) was only present in RIH samples and the phylum Euryarchaeota (Archaea) in NDRI samples.

**Figure 1.**
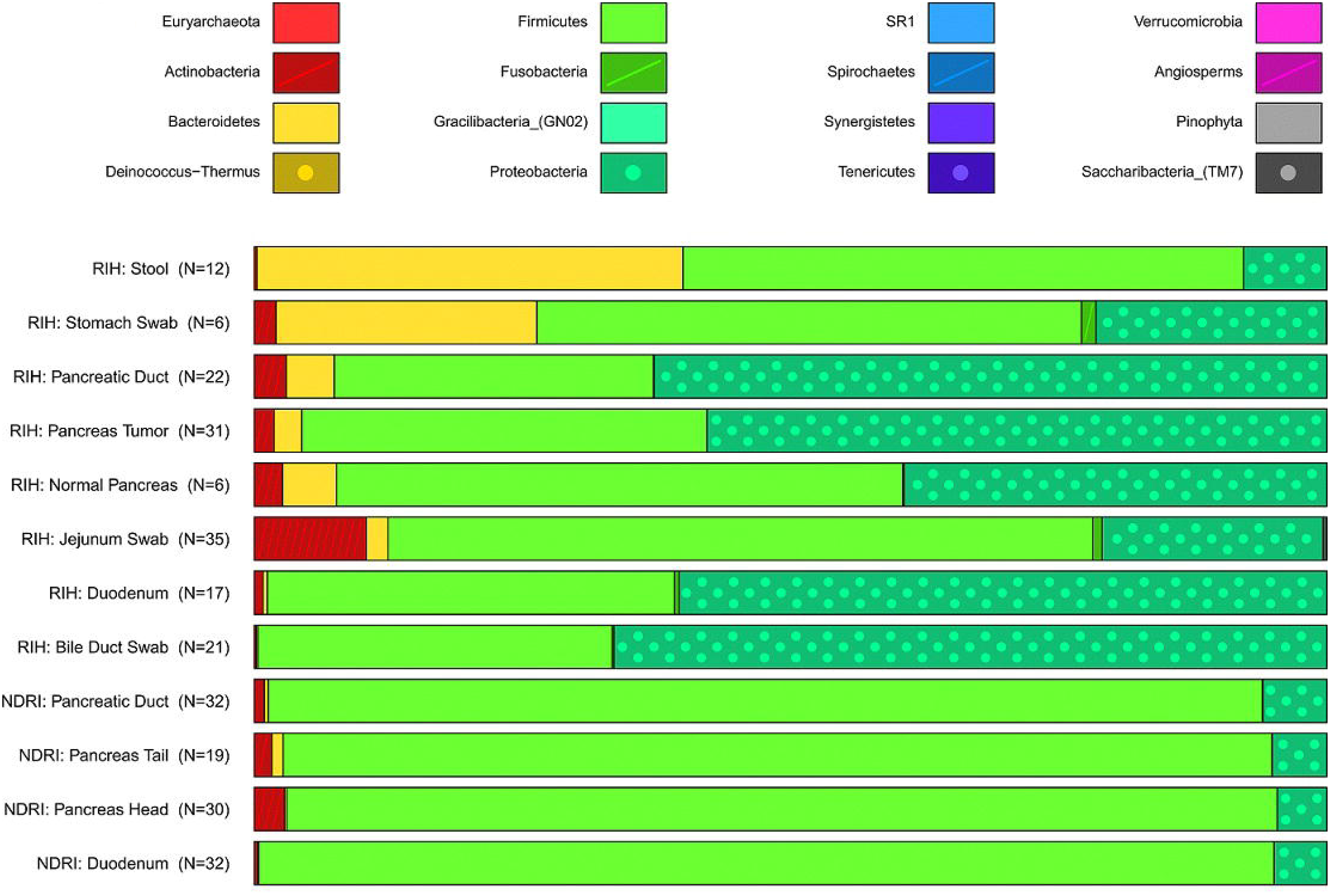
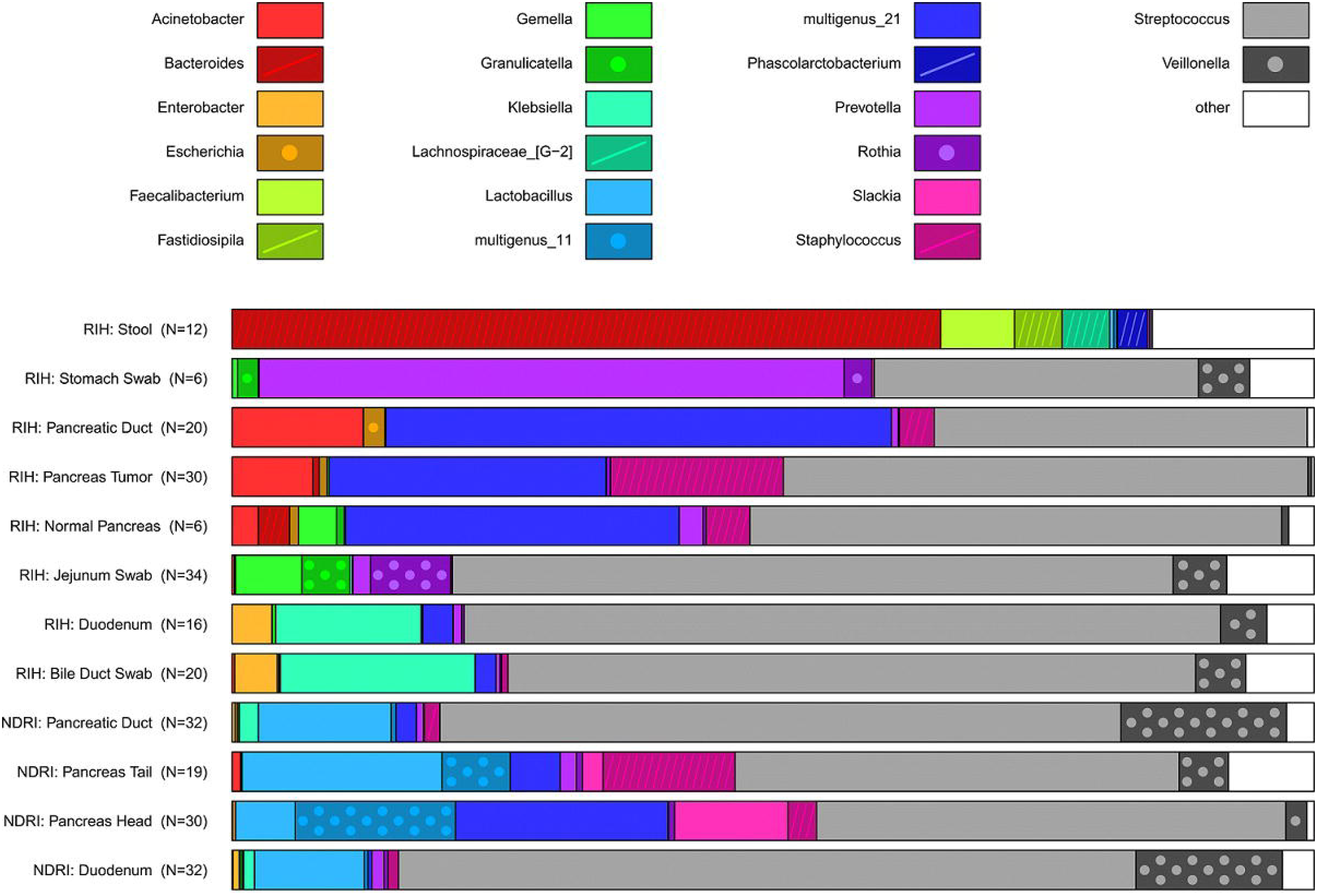
Distribution of bacteria relative abundance by (a) phylum level and (b) genus level in all the studied body habitats based on read taxa attribution using V3-V4 hypervariable region of 16S rRNA genes.

While the microbial communities in the pancreatic tissues were dominated by the phyla Firmicutes and Proteobacteria, substantial inter-individual variability was observed. In RIH samples, Proteobacteria relative abundance ranged from 2 to 99%, and similarly, Firmicutes relative abundance ranged from 0.6 to 84%. Large inter-individual variability was also observed in the NDRI samples. Proteobacteria primarily consisted of the family Enterobacteriaceae (genera *Enterobacter* and *Klebisella*) and unclassified gammaproteobacteria.

Median relative abundance at the phyla (Figure 1A) and genus taxonomic levels (Figure 1B) were similar for tissue samples obtained from the pancreatic tumors and pancreatic ducts in RIH patients, but differed from those in the duodenum, bile duct and jejunum. Bacteria from the genera *Klebsiella* and *Enterobacter* were more commonly found in the duodenum (RIH and NDRI samples) and bile duct samples (RIH) (Figure 1B), and bacteria from the genera *Gemella, Granulicatella, Prevotella* and *Rothia* were more common in the jejunum samples (Figure 1B), compared to the pancreatic samples. High levels of Bacteriodetes were noted in the stool samples and stomach swabs (Figure 1A); the majority of these bacteria belonged to the genus *Bacteroides* in the stool and *Prevotella* in the stomach (Figure 1B). All samples, except the stool samples, had high levels of *Streptococcus* (Figure 1B). Pancreatic tissue samples from the NDRI subjects had higher levels of *Lactobacillus* and lower levels of *Acinetobacter* than those from the RIH (Figure 1B).

### Within and between sample diversity analysis

With the exception of the stool and the jejunum, all the bacterial communities were characterized as habitats with low bacterial richness including the pancreatic sites, duodenums and the bile ducts (Figure 2A). Among RIH patients, the microbial communities of the stool samples were represented by higher richness than the microbial communities in the tumors of the pancreas (*p*=0.007), duodenums (*p*=0.013) and bile duct swabs (*p*=0.017). Likewise, the stool bacterial communities had higher richness than the NDRI pancreatic heads (*p=*0.012), pancreatic ducts (*p*=0.020) and duodenums (*p=*0.005). The microbial communities in the jejunum swabs from the RIH samples showed more richness than the communities in the RIH pancreatic head (*p=*0.014) and duodenums (*p=*0.028). In general, the bacterial communities in the pancreas of RIH patients were similar in richness when compared to those from the pancreas of the NDRI matching sample types. Similar results were observed using additional alpha diversity measures of the bacterial communities (Figure 2B-D). As expected, the stool samples were the most diverse with a Shannon index **≥** 4 ^24^(Figure 2B). As the number of phyla represented in high abundance in these samples was relatively low (∼5), we observed relatively low levels of phylogenetic distances across all samples (Figure 2D).

**Figure 2.**
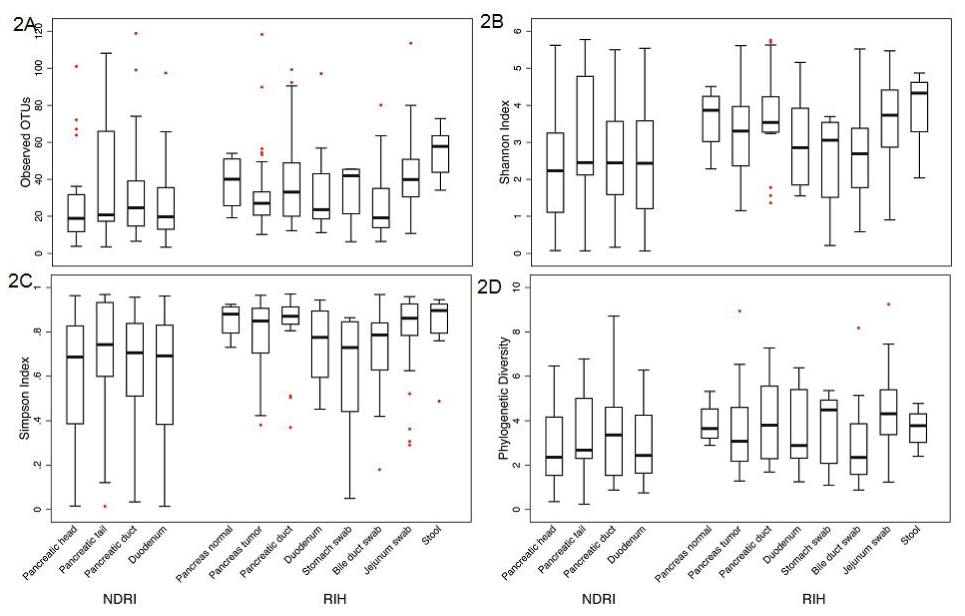
Comparative alpha diversity analyses of bacterial communities in anatomical sites (based on a simulated data set subsampled from the input OTU table). Alpha diversity metrics: (A) Richness, (B) Shannon diversity index, (C) Simpson index, and (D) Phylogenetic diversity.

The ordination beta-diversity analysis revealed that the majority of samples belonged to a single cluster, without any apparent grouping by health status or by anatomical site; the only exception were the microbial communities from the stool samples and the majority of the jejunum swab samples that formed two additional distinct clusters (Figure 3). However, significant differences in microbial composition were noted between duodenums from RIH and NDRI subjects (*p*=0.03), and between the bile-duct, stomach, and jejunum swabs (*p*=0.0002).

A β-diversity analysis using the Euclidean distance to describe the relatedness of microbial communities across anatomical sites further confirmed that samples from the same subject were significantly more related to one another than samples from the same anatomical site in different subjects (Figure 4).

**Figure 3.**
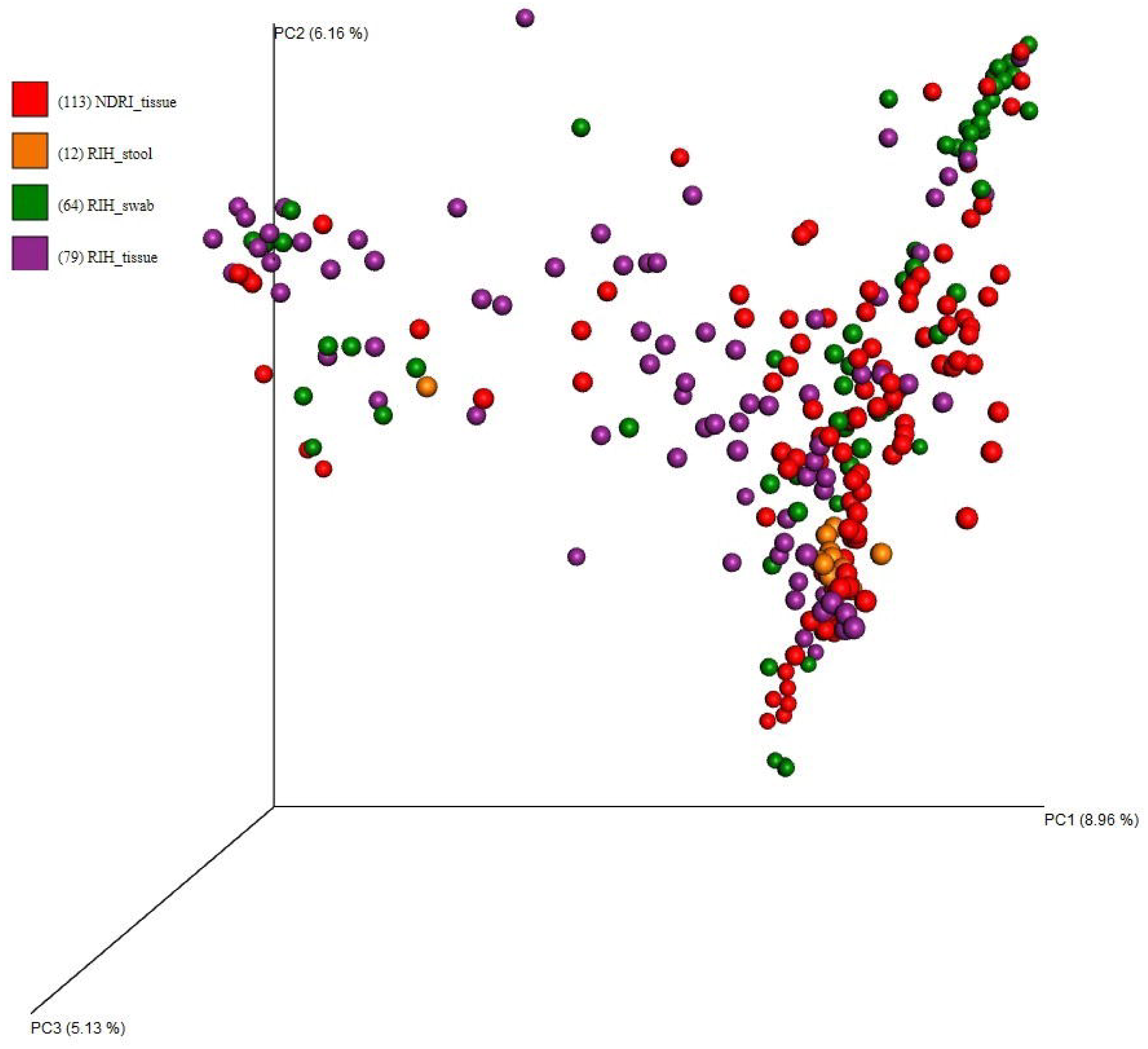
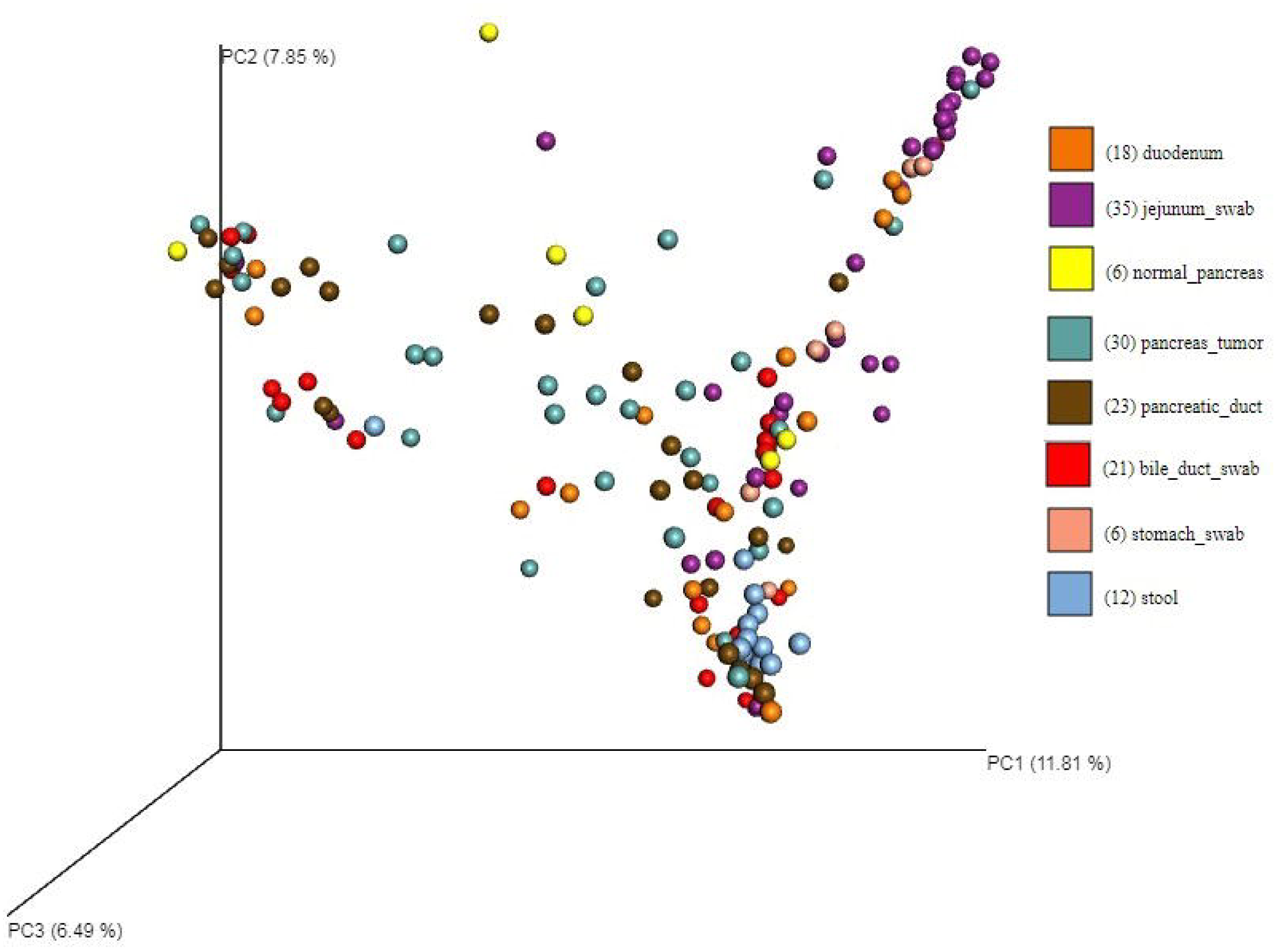
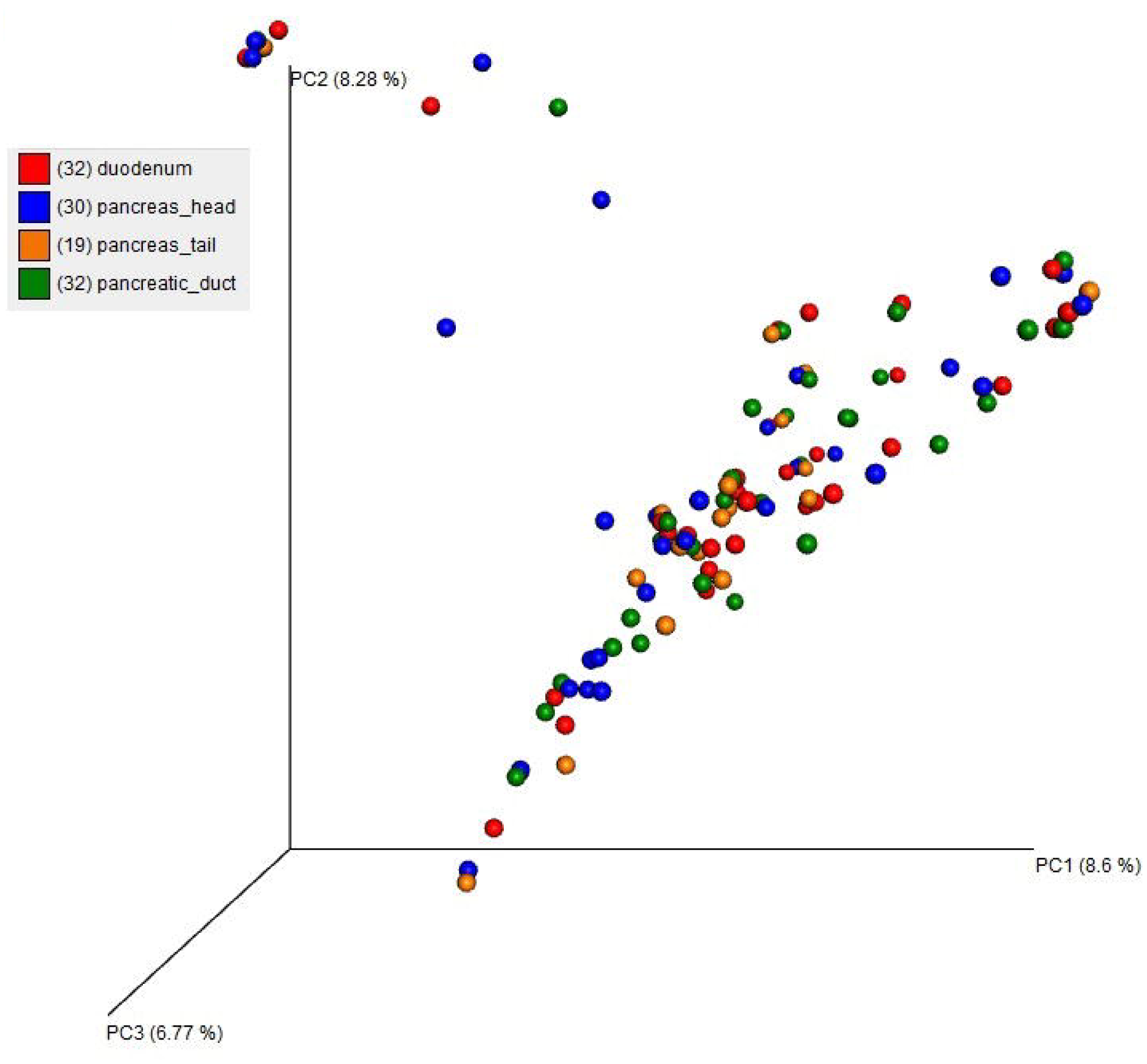
PCoA plots showing the relatedness of microbial communities among samples from RIH patients and NDRI donors using the Bray-Curtis dissimilarity index. Individual datasets are colored according to their (A) RIH and NDRI sample type, (B) RIH anatomical site, and (C) NDRI anatomical site.

**Figure 4.**
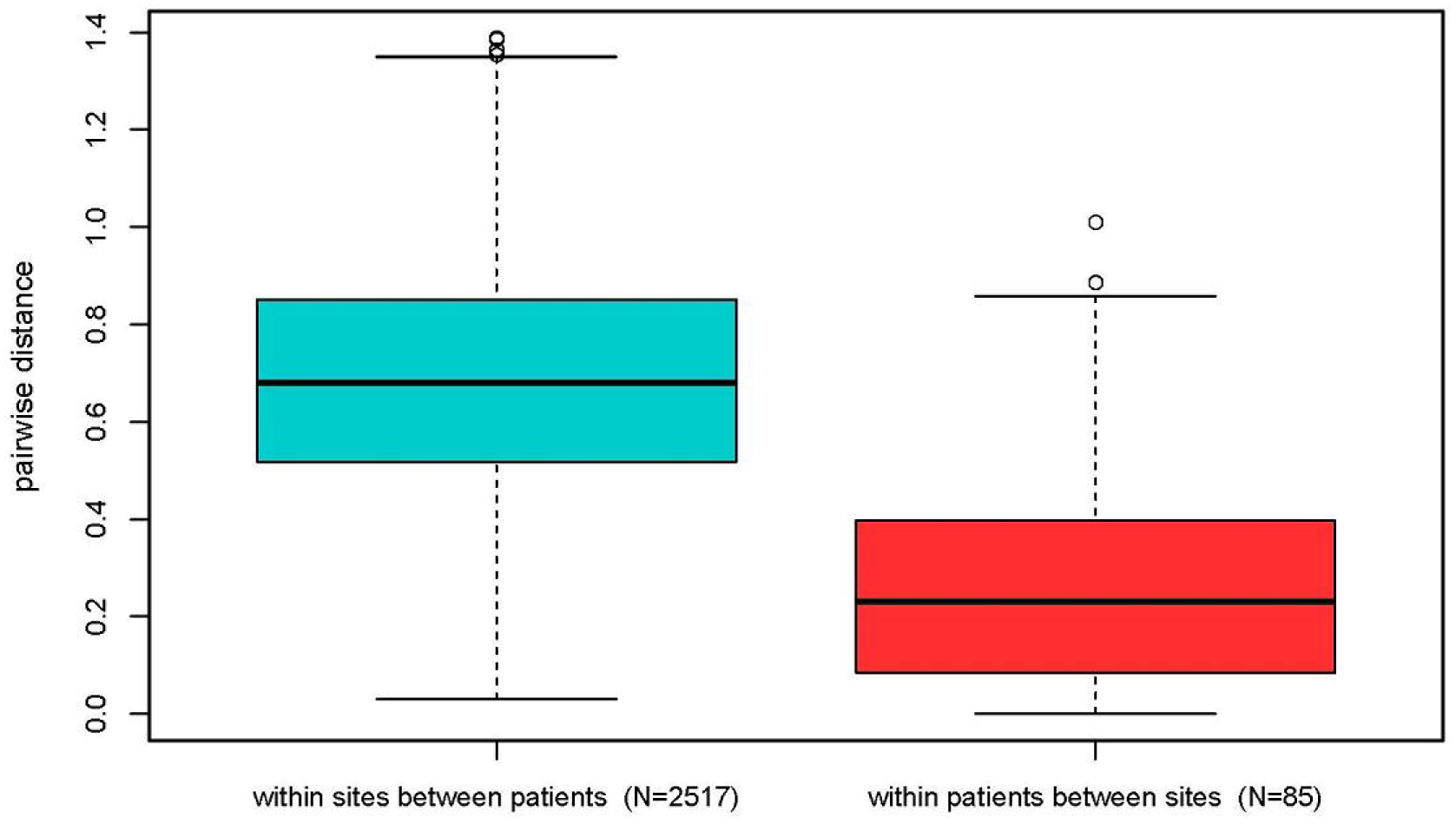
Euclidean Distances based on relative abundance within sites between subjects, and within subjects between sites for RIH and NDRI groups.

### Associations of host factors with microbial communities

We explored which factors obtained in the questionnaires and medical files in RIH patients were associated with bacterial communities in the pancreatic tissue samples. The most influential factors were sequencing run, presence of stent, and chemotherapy prior to surgery; each of these factors was significantly associated with a large number of genera tested in marginal models. Considering these factors may confound the main associations of interest, we included them in the models comparing the different ICD-codes among the RIH patients. Age, sex, BMI, and smoking status were not associated with the majority of the genera tested; however, age, sex and BMI were kept in the models to be conservative.

Using multiple regression analyses, we examined presence or absence, and relative mean abundance of bacterial taxa (at the genus level) in pancreatic tissue by disease phenotype (tumors/cysts) compared to pancreatic tissue (pancreatic head) from the NDRI samples. Table 2 presents the bacterial taxa for which statistically significant associations remained after Bonferroni correction with adjustment for age, sex, BMI and run; each model includes both a zero-inflated component and a relative mean abundance comparison. In the marginal models (prior to adjusting for other covariates), a total of 25 genera were identified as being significantly associated with disease status prior to correction for multiple comparisons (Supplemental Table 2). Of the original 25 genera, 7 are known oral bacteria that have been previously implicated in periodontal diseases (*Porphyromonas, Pyramidobacter, Aggregatibacter, Capnocyophaga, Prevotella, Selenomonas,* and *Gemella*). Mean relative abundances for these taxa (namely, *Pyramidobacter, Capnocyophaga, Prevotella, Selenomonas,* and *Gemella)* were higher in samples coming from patients diagnosed with pancreatic cancer (ICD C25) and/or presence of bacterial taxa was more common in pancreatic cancer (*Haemophilus, Pyramidobacter, Aggregatibacter, Prevotella,* and *Gemella*). The model with *Porphyromonas* had the strongest association overall (*p*=1.2x 10^−7^); the relative mean abundance for periampullary cancer tissue samples was substantially higher than that of NDRI samples (*p*=5.8 x 10^−19^), as were the IPMNs (K86.2) samples (*p*=3.6 x 10^−7^). The associations with *Porphyromonas* remained elevated in multiple regression models (Figure 5; Table 2).

**Figure 5.**
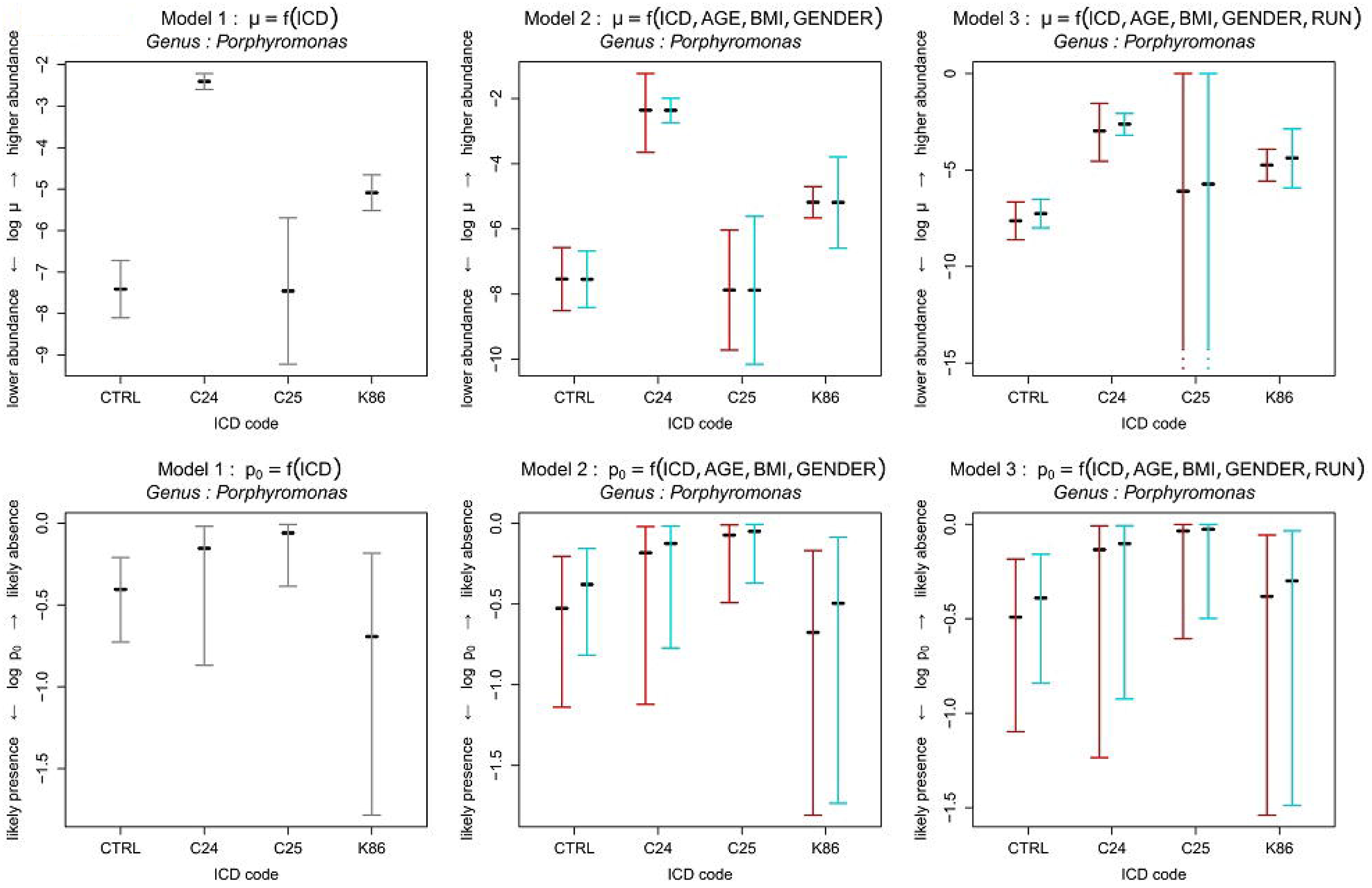
Plots of results (log mean and log probability of absence vs presence) for genus *Porphyromonas* obtained from zero-inflated regression models including NDRI pancreatic tissue samples (head of pancreas) and RIH pancreatic tissue samples (by ICD code). Three models are presented: the marginal model (Model 1); the model adjusting for age, BMI, gender (Model 2); and the model adjusting for age, BMI, gender, and sequencing run (Model 3). (Red = Females and Blue = Males)

**Table 2.**
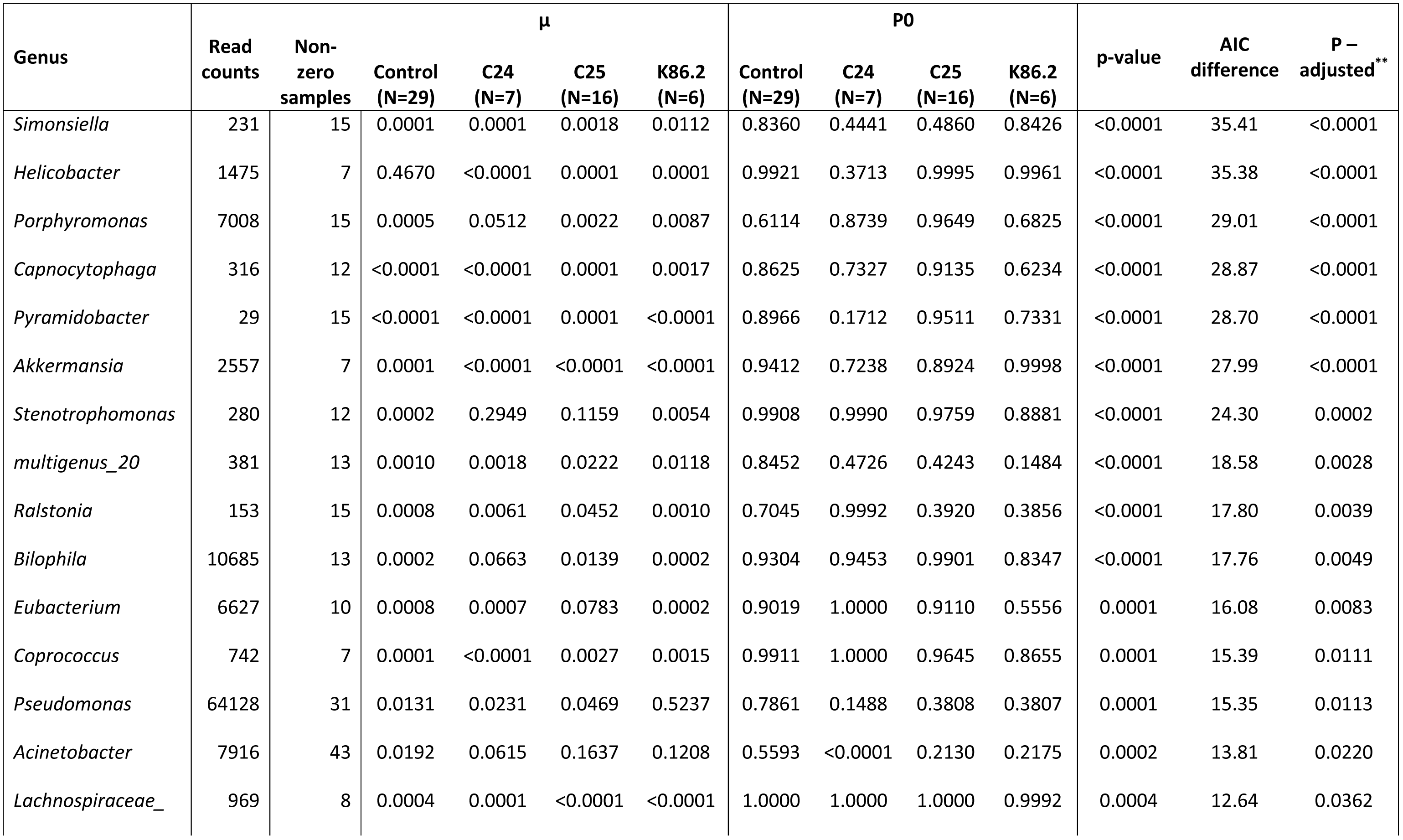
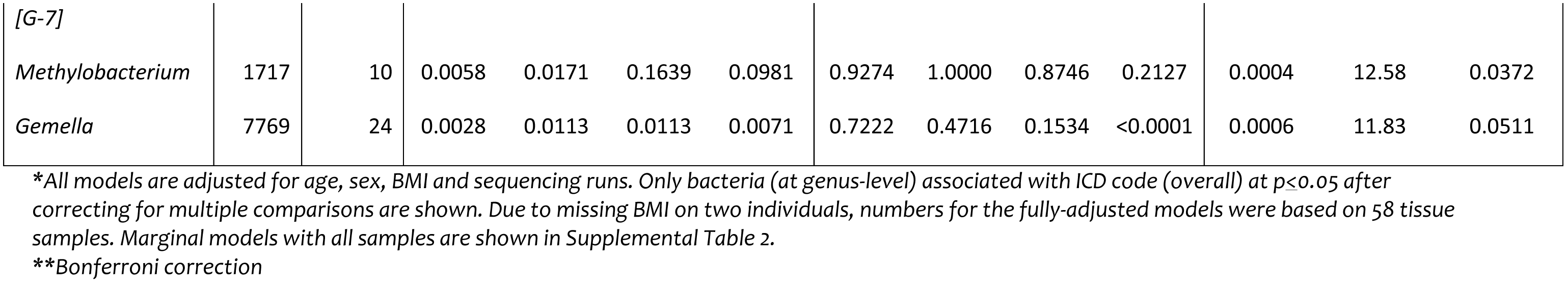
Multiple regression models including zero-inflation parameter comparing bacteria in pancreatic tissue from NDRI and RIH samples*

In the marginal model, the second strongest association was observed for the genus *Pyramidobacter*; this bacteria, for which only one species is known, was more likely to be identified in periampullary pancreatic cancer (71%) and IPMNs (33%), than in NDRI pancreatic head samples (23%); however, count numbers were low in all samples (Table 2). The multivariable regression models for the pancreatic tissue samples identified 13 bacterial taxa (at the genus-level) that had not been significant in the marginal regression models, including *Simonsiella, Helicobacter, Akkermansia,* and *Bilophia* (Table 2 vs Supplemental Table 2). *Helicobacter* was commonly identified in periampullary pancreatic tumors (C24) but at low levels; in contrast, *Helicobacter* was infrequently identified in the NDRI samples, but was a dominant genus when present (relative mean abundance 47%; Table 2).

We further examined the RIH pancreatic tumor tissue samples without the NDRI samples given the difference in source of tissue. In these models, comparing the different ICD codes, we were able to adjust for potential confounding by chemotherapy and stent. *Porphyromonas* and *Pyramidobacter* were also strongly associated with ICD code in the marginal models (Supplemental Table 3) and multiple regression models.

To test whether the associations would be similar using pancreatic **duct** tissue samples, we repeated the analysis using RIH and NDRI samples obtained from the pancreatic ducts (non-tumor tissue for pancreatic cancer patients). The associations for *Porphyromonas* remained detectable and statistically significant in these analyses (*p*=1.53 x 10^−11^), and another oral bacterial taxon associated with periodontal disease (e.g., *Tannerella*) emerged in these samples as being more abundant in the periampullary pancreatic cancer patients (C24) in the marginal model, but was not statistically significant in the multiple regression model.

Using tissue samples obtained from the **duodenum**, we compared relative abundance of bacteria in NDRI and RIH patients to examine whether any bacteria from the pancreatic tissue analyses were also noticeably different in the duodenum samples. Of the significant associations noted in the pancreatic tissues, *Selenomonas* was also elevated in the duodenum tissue of pancreatic cancer patients (*p*=3.9 x 10^−12^) compared to duodenum tissue from NDRI patients. A weak association was also observed for *Gemella* for the duodenum samples, consistent with an overall elevated mean relative abundance in the RIH samples compared to the NDRI samples; other associations were either not significant or not consistent in direction of differences.

We only analyzed the microbial community of one duodenal stent. The bacteria present were characterized as the members of the genera *Klebsiella* and *Enterobacter*.

## Discussion

Using pancreatic tissue samples from patients with pancreatic cysts or pancreatic cancer, and comparing them to pancreatic tissue samples obtained from donors who died of non-cancer causes, we were able to demonstrate for the first time that pancreatic tissue contains a number of different bacterial taxa, including taxa that are known to inhabit the oral cavity. To our knowledge, our metagenomics targeted analysis, using the 16S rRNA bacterial gene, was the first to be conducted on fresh pancreatic tissue samples. Our findings provide evidence that the pancreas is not a sterile organ and that there is substantial inter-individual variability in relative abundance of different bacteria at the genera and phyla level. In addition, we show that the bacterial composition at different sites in the pancreas (i.e., duct, head and tail) are similar in the same individuals. Finally, we noted significant differences in abundance of periodontal-related pathogens in the tissue of pancreatic patients when compared to non-cancer patients.

Previous studies have reported associations between periodontal disease pathogens and pancreatic cancer risk, especially *Porphyromonas gingivalis* ^6 7^. Periodontal disease is an inflammatory disease of the gums that can, in advanced conditions of periodontitis, result in systemic inflammation. Dissemination oral bacteria has also been observed and linked to a number of chronic diseases, including cardiovascular diseases ^25 26^. *Fusobacterium nucleatum* has been associated with colon cancer in several cross-sectional studies ^27 28^. However, it remains unclear if the bacteria identified in those studies are opportunistic or causal ^29^; no prospective study on colon cancer has observed associations with this bacterium prior to cancer onset, although a number of studies have reported positive associations between periodontal disease and colon cancer risk ^30^. Mouse models of colorectal cancer provide some support for a causal link ^31^, demonstrating how this bacterium has the ability to initiate recruitment of tumor-infiltrating immune cells.

In this study, we observed significantly higher mean relative abundance levels (at the genus-level) of a number of bacterial taxa previously associated with severe periodontitis in pancreatic tissue, including *Porphyromonas, Aggregatibacter, Prevotella, Gemella* and *Selenomonas* ^32-34^; however, only *Porphyromonas* and *Gemella* remained statistically significant after adjusting for age, sex, BMI and sequencing run*. Porphyromonas* was also elevated in the pancreatic duct tissue of periampullary pancreatic cancers, but no statistically significant associations were noted for the other bacterial taxa. Mean relative abundance was also higher for *Gemella* when comparing duodenum tissue from pancreatic cancer patients to NDRI duodenum samples. Whether any of these bacteria play a role in pancreatic carcinogenesis will need to be further examined in other studies and confirmed in animal models. Proposed mechanisms for carcinogenesis include the ability of certain bacteria to induce a pro-inflammatory response in the tumor microenvironment ^31^; inhibit the immune response targeted at eliminating tumor cells ^35^; and modulate key cellular pathways associated with cell division ^36^.

A similar study using swab specimens from the pancreas, bile and jejenum, was conducted on patients with pancreatic cancer undergoing pancreaticoduodenectomy ^13^. In that study, a large number of bacteria were present in fluids obtained from the pancreatic ducts and the common bile duct, including *Prevotella, Haemophilus, Aggregatibacter,* and *Fusobacterium* ^13^. Consistent with our findings, microbial communities in the pancreas, bile and jejunum fluids were similar within individuals ^13^. Mean relative abundance for the bacterial taxa *Klebsiella* was high in the samples from pancreatic cancer patients in that study ^13^; in our study we found this bacteria to be one of two present on a swab taken of the stent itself. Placement of stent prior to surgery may impact the type of bacteria present in the pancreas, as observed in our study. In a separate study, metagenomics was conducted on fresh frozen duodenum samples from 5 normal and 5 obese individuals; *Streptoccocus* (30-32%) and *Actinomyces* (12-17%) were the most common bacterial taxa identified in those samples, and relatively higher counts of *Gemella* were also identified in all 10 subjects ^37^. *Porphyromonas* bacterial taxa were not identified in the duodenal samples ^37^.

Several studies have looked at the involvement of bacteria in biliary and pancreatic diseases and have observed a high number of bacterial taxa present in the calcified pancreatic duct epithelium and in pancreatic abscess ^8 38-41^. Anaerobic bacterial taxa have been found at a variable rate in pancreatitis; the results depend on the process for bacterial identification ^8 38 39^. Previous studies have also reported the presence of bacteria in bile ^42 43^. In a study of 6 patients with gallstones, 16S rRNA gene sequencing identified high relative abundances of *Escherichia, Klebsiella* and *Pyramidobacter* in the bile, and the bacterial profile of the bile was very similar to the duodenum in the same patients ^43^. In our study, mean relative abundance of *Pyramidobacter* in pancreatic tissue was strongly associated with pancreatic cancer, although levels of read were low in these samples. This species was originally isolated from the oral cavity ^44^. Given that gallstones have been associated with risk of pancreatic cancer in some studies ^45^, the role of this bacterium should be further examined.

A number of the bacterial taxa we observed with elevated relative mean abundance in RIH samples have been previously identified in immunocompromised patients and are largely believed to be opportunistic pathogens, including *Acinetobacter* ^46^, *Kluyvera* ^47^, and *Stenotrophomonas* ^48^. The genus *Gemella*, which was found at higher relative abundance in pancreatic cancer patients when compared to NDRI samples in both pancreatic tissue samples and duodenum tissues, has been previously associated with endocarditis ^49 50^. Because our analysis was based on a cross-sectional study design, we expected to identify bacteria that were present as a result of opportunistic nosocomial infections given that the majority of RIH patients were likely immunocompromised from their cancer. However, our results show that even normal pancreatic tissue harbor a microbial community.

The strength of this study was the collection of specimens specifically for the purpose of microbiome analysis, with numerous precautions made to reduce contamination during collection and processing of samples. Moreover, multiple types of samples were collected on each patient at RIH, including obtaining tissue or swabs from multiple sites, to allow for inter vs. intra-individual differences at different sites. Finally, the multivariable regression analyses was conducted to adjust for potential confounding by known pancreatic cancer risk factors, including BMI and smoking, as well as other factors that may cause bias, including pre-OP EUS and prior chemotherapy.

The major limitation of this analysis was the small number of patients with pancreatic cysts and pancreatic cancer; despite recruiting 77 patients, not all subjects had tissue resections during surgery (as more advanced pancreatic cancer patients are often not operable) and our analysis was based on a subsample of that population with tumor samples (n=30). Furthermore, there appeared to be differences in bacterial composition between periampullary pancreatic cancer patients (ICD 24) as compared to those with pancreatic ductal adenocarcinoma (ICD 25), which further reduced our sample size. More research is needed to explore the differences in bacterial profiles between precursor lesions (i.e., IPMNs) and site of cancer origin.

In this study, we detected a number of bacterial taxa in pancreatic tissue, from cancer patients as well as non-cancer subjects and demonstrate that the pancreas has a microbiome distinct from the gut (jejunum) and stomach. Furthermore, the bacterial profiles in the pancreas were more similar within individuals across different sites of the pancreas (i.e., head, tail, ducts) than between individuals at each site. Bacterial taxa known to inhabit the oral cavity were common in the pancreas microbiome and periodontal pathogens were also identified in many pancreatic tissue samples. Despite small numbers, we were able to identify higher relative abundance of *Porphyromonas* in pancreatic tissue samples of cancer patients compare to those samples obtained from the NDRI, and associations remained statistically significant after adjusting for a number of potential confounders and for multiple comparisons. Further research is needed to address if and how these bacterial taxa may be related to carcinogenesis.

## Acknowledgement

We thank the participants who graciously enrolled in this study. We thank Ms. Priyanka Joshi for her tremendous help with recruitment of patients at the RIH, and Dr. Ross Taliano for assisting with the preparation of the tissue specimens in the Pathology Department at the RIH. We also thank Drs. Murray Resnick, Kara Lombardo, Emily Walsh and Laura Gantt for their help with this project.

